# An Efficient Method to Quantify Structural Distributions in Heterogeneous cryo-EM Datasets

**DOI:** 10.1101/2021.05.27.446075

**Authors:** Hanlin Gu, Wei Wang, Ilona Christy Unarta, Wenqi Zeng, Fu Kit Sheong, Peter Pak-Hang Cheung, Song Liu, Yuan Yao, Xuhui Huang

## Abstract

Cryogenic Electron Microscopy (cryo-EM) preserves the ensemble of protein conformations in solution and thus provide a promising way to characterize conformational changes underlying protein functions. However, it remains challenging for existing software to elucidate distributions of multiple conformations from a heterogeneous cryo-EM dataset. We developed a new algorithm: Linear Combinations of Template Conformations (LCTC) to obtain distributions of multiple conformations from cryo-EM datasets. LCTC assigns 2D images to the template 3D structures obtained by Multi-body Refinement of RELION via a novel two-stage matching algorithm. Specifically, an initial rapid assignment of experimental 2D images to template 2D images was applied based on auto-correlation functions of image contours that can efficiently remove the majority of irrelevant 2D images. This is followed by pixel-pixel matching of images with fewer number of 2D images, which can accurately assign the 2D images to the template images. We validate the LCTC method by demonstrating that it can accurately reproduce the distributions of 3 *Thermus aquaticus* (*Taq*) RNA polymerase (RNAP) structures with different degrees of clamp opening from a simulated cryo-EM dataset, in which the correct distributions are known. For this dataset, we also show that LCTC greatly outperforms clustering-based Manifold Embedding and Maximum Likelihood-based Multi-body Refinement algorithms in terms of reproducing the structural distributions. Lastly, we also successfully applied LCTC to reveal the populations of various clamp-opening conformations from an experimental *Escherichia coli* RNAP cryo-EM dataset. Source code is available at https://github.com/ghl1995/LCTC.

## 1 Introduction

Cryogenic Electron Microscopy (cryo-EM) [1] has become one of the most popular techniques to elucidate the structures of biomolecules at atomic resolution [2, 3]. Different from X-ray crystallography, cryo-EM has the advantage of capturing an ensemble of biomolecule conformations. Cryo-EM achieves this by flash-freezing the sample solution, which preserves the heterogeneous conformations of the biomolecules in the solution state. On the other hand, X-ray crystallography typically requires a large amount of homogeneous samples [4], rendering it unsuitable for the analysis of biomolecular conformational changes. Also, cryo-EM requires a much smaller amount of samples and accepts a larger variation of specimen types, including single protein molecules, large protein complexes, thin-protein crystals, virus particles, helical fiber complexes, bacteria, cells, and even entire tissue sections [5].

Many biomolecules are intrinsically flexible and undergo large conformational changes to perform their function. Cryo-EM becomes an important technique to identify various 3D conformations of biomolecules, since it collects thousands of 2D images that span the biomolecular conformations in 3D space. The flash-freezing technique in cryo-EM allows the capture of an ensemble of biomolecule conformations that approximates the solution state following the Boltzmann distribution. Thus, the distribution of the conformations naturally reflects the underlying free energy landscape of the biomolecule. Therefore, cryo-EM technology is useful to study macromolecules in their native conditions, in which multiple metastable conformations may co-exist [6, 7]. Furthermore, investigating the population distribution of the conformations in the cryo-EM dataset can help us understand the thermodynamics of the macromolecules.

However, the classification of heterogeneous data in the single-particle reconstruction of biological molecules into several conformational classes has two major challenges. Firstly, conformational changes underlying protein function are often localized and thus difficult to be detected. Secondly, conformational changes can be obscured by the viewing angles.

There are mainly two types of methods to address the above-mentioned challenges. The first type is based on the Maximum Likelihood to reconstruct a given number of structures from one cryo-EM dataset, which sets up a probabilistic model with respect to the model parameters given to the cryo-EM 2D images. Popular software such as RELION [8, 9] and cryoSPARC [10] are based on Maximum Likelihood approach, in which a statistical noise model is employed to calculate posterior probabilities for all possible orientations and classes. For example, Nakane et al. [11] applied Multi-body Refinement analysis in RELION to investigate the molecular motions of a structural feature (a region or a domain) in the biomolecule. In addition, an algorithm based on cryoSPARC called “3DVA” was recently developed [12], where the conformational landscape of a molecule was modelled in a linear subspace as superposition of conformations weighted by their probability, and then fitted into a high resolution volume. All these Maximum Likelihood-based methods estimate the classifications based on the reconstructed 3D volumes. However, all of them cannot informatively elucidate the distributions of different conformations.

Another type of method is to directly cluster the experimental 2D-images. One idea is to use a specific type of manifold learning for dimensionality reduction of the 2D images to analyze heterogeneous cryo-EM datasets [13]. Pencezk et al. employed Principal Component Analysis (PCA) of cryo-EM data and performed clustering to assign the 2D images into different conformations. Other methods treat the manifold as a complex and nonlinear geometry and the diffusion map was used to search for the mapping of heterogeneity [14, 15]. In addition, Shatsky et al. developed a spectral clustering method that is based on image similarity of “common line” [16]. However, these direct clustering methods impose a common difficulty. For instance, the projection of 3D shapes onto a few collective variables or manifolds may obscure the functional conformational changes of interest. Therefore, the development of an automatic and efficient algorithm to elucidate multiple metastable conformations and their equilibrium populations from a single heterogeneous cryo-EM dataset is needed.

In this work, we developed the Linear Combinations of Template Conformations (LCTC) algorithm to accurately and efficiently obtain the populations of multiple conformations in a single cryo-EM dataset. We first determined a few template 3D conformations of a biomolecule using Multi-body Refinement in RELION [11]. To improve the accuracy of our classification method, we minimized the bias due to the projection direction (or viewing angle) by identifying a viewing angle that can best distinguish the conformational changes of interest. We then projected each template 3D conformation onto 2D images at the chosen viewing angle as the template 2D-images. Next, 2D images from the experimental cryo-EM dataset were assigned to template 2D images via an efficient two-stage matching method. In the first stage, a fast initial clustering was performed based on the rotational invariant contours of the 2D image. This is followed by the second stage, in which a more accurate pixel-pixel based image matching was conducted. After the two-stage matching process, we could obtain the populations of multiple conformations in a single cryo-EM dataset. We have demonstrated that our approach could be applied to elucidate the conformational distributions efficiently and more accurately than the Clustering-based and Maximum Likelihood-based methods in two systems of the dataset. For the opening of the clamp domain of *Thermus aquaticus* (*Taq*) RNAP, we can reproduce the population of RNAP in a simulated cryo-EM dataset. Furthermore, we have successfully applied the LCTC method to an experimental dataset describing the Clamp domain motion of *Escherichia coli* (*Eco*) RNAP.

## 2 Methods

Our scheme is composed of four steps: 1) The best viewing angle to differentiate conformational changes in template images is identified. 2) Denoising cryo-EM 2D images is performed and the template 3D structures, generated by multi-body Refinement of RELION, are projected around the best viewing angle, which generate template 2D images, to prepare for matching step. 3) cryo-EM 2D images are assigned to template 2D images via a two-stage matching process, in which the first matching is based on novel Euclidean distance based on Auto-correlation Functions (ACF) followed by matching based on the accurate Euclidean distance calculation of pixel to pixel (pixel-pixel). 4) Lastly, we count the number of cryo-EM 2D images assigned to each template conformation, which represents the distribution of conformations in cryo-EM.

### 2.1 Identification of the best viewing angle

Biomolecules have complicated 3D structures, and their functional conformational changes often involve just a subset of the system. The first step of our protocol is to identify the best viewing angle that can best distinguish the conformational changes. We define the optimal viewing angle as onto which the projections of various template 3D conformations produce the largest variations. As an example: we plot the projection of three 3D conformations of *Taq* RNAP (Figure 2 (a)-(c)) on the best viewing angle (Figure 2 (d)-(f)) and a bad viewing angle (Figure 2 (g)-(i)). Clearly, the projection at the best viewing angle can distinguish the three conformations, while the bad viewing angle cannot.

**Figure 1:**
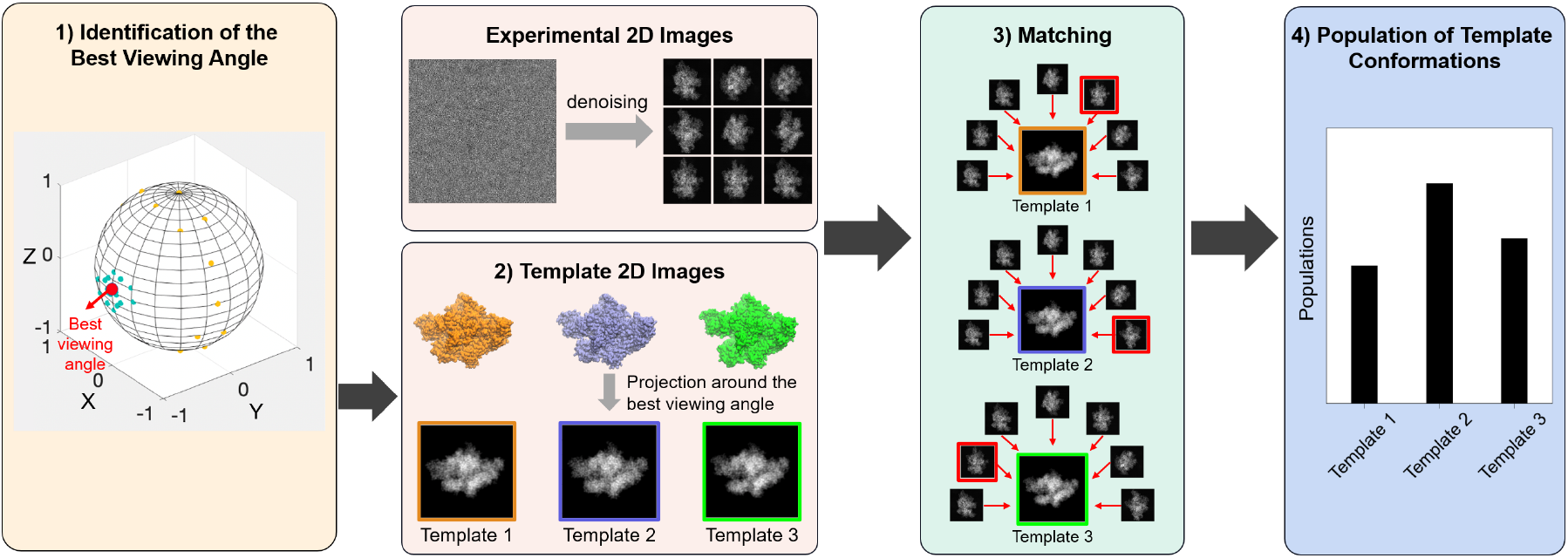
A schematic overview of LCTC. The procedure of LCTC method is summarized as follows, 1) the best viewing angle, that can best differentiate the conformational changes of the biomolecule, is identified using the template 3D conformations, 2) the template 3D conformations are projected around the best viewing angle to generate template 2D images, 3) the experimental cryo-EM 2D images are assigned to the template 2D images using the two-stage matching, 4) the number of cryo-EM 2D images assigned to each template image is counted and the populations of the template conformations are estimated.

**Figure 2:**
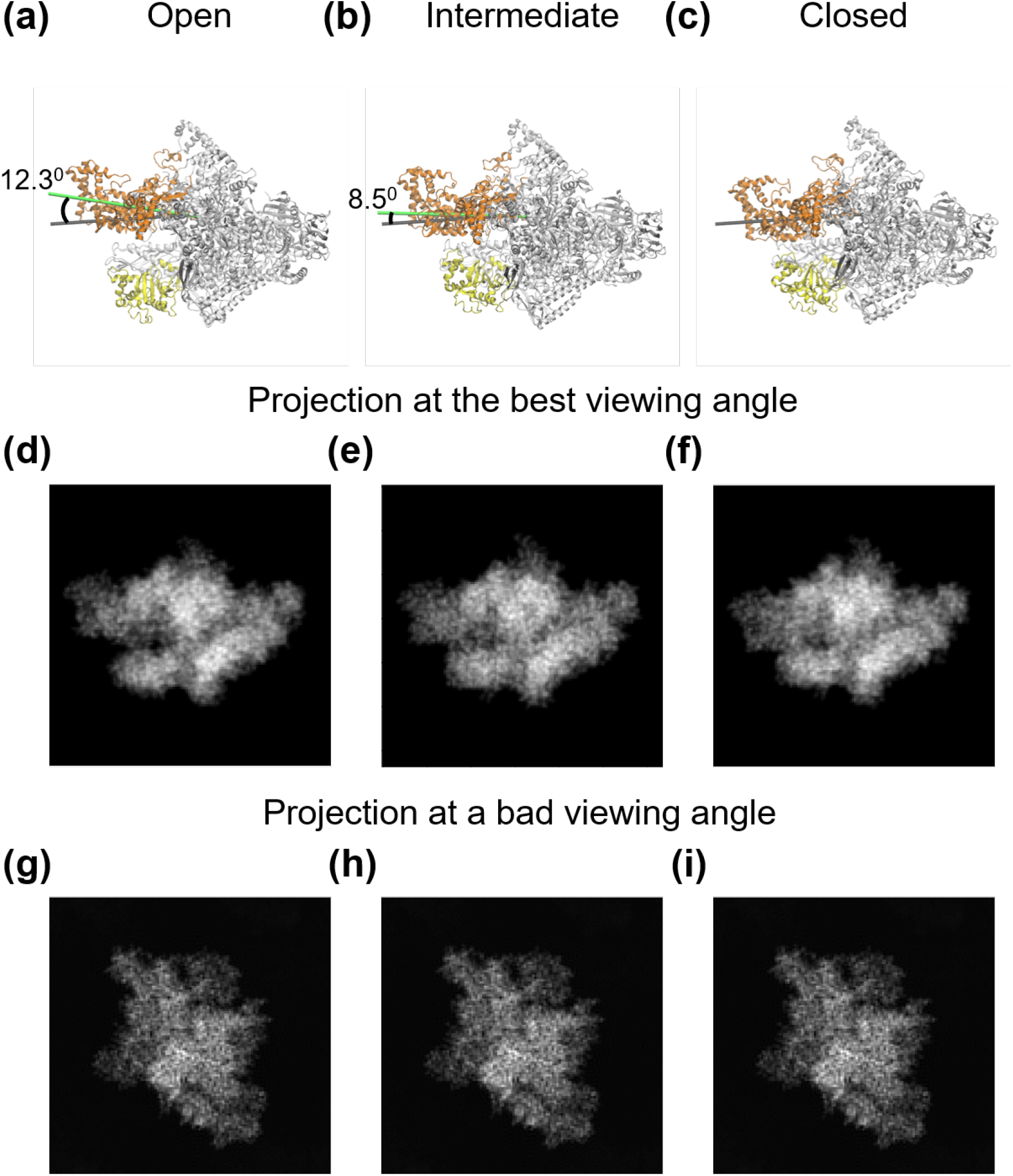
Identification of the best viewing angle. The best viewing angle is the angle that can best distinguish the conformational changes of the biomolecule. In this example, the 3D volumes of three template *Taq* RNAP conformations, (a) Open, (b) Intermediate, (c) Closed were projected onto the 2D images (d), (e), (f) at the best viewing angle. The same 3D volumes were also projected at a bad viewing angle (g), (h), (i).

We firstly obtained *K* representative conformations (template structures) from the Multibody Refinement to generate the corresponding 3D density maps. We denoted *A* as a *K* × *N*^3^ size matrix consisting of *K* conformations *A_i_* (*i* = 1, 2, …*K*) and *N*^3^ as the dimension of 3D density maps, *P* = *R_z_* · *R_y_* · *R_z_* · *P*_0_ is a projection in ℝ^3^ which is the product of rotational matrix *R_z_*, *R_y_*, *R_z_* representing the viewing angles in discretized sphere space Ω, and the initial projection *P*_0_ representing the projections from the north pole. Our aim is to maximize the variance of projections of *K* conformations: Var(*PA*_*i*_, *i* = 1, 2, …, *K*), where *PA*_*i*_ denotes the projection of discrete 3D density map *A_i_*.

After performing PCA on *A*, we obtained the eigenvalues λ*_j_* and eigenvectors *V_j_* of covariance matrix Σ = *A^T^ A*. We approximated the variance of projected 3D density maps based on the eigenvalues and eigenvectors as shown in Equation (1). In other words, projections of top *T* eigenvectors 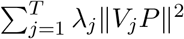 could be applied to directly estimate the variance of projections of *A* as Var(*P A_i_*, *i* = 1, 2, …*K*) (see Appendix 6.1 for the proof).

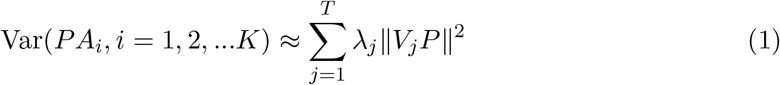

With top *T* eigenvectors {*V_1_*, …, *V_T_*}, we then performed eigenvector projections using the projection matrix *P* of different viewing angles and find the one that produces the largest variance to get the best projection matrix *P*\* as Equation(2) showed. After obtaining the best viewing angle in *P*\*, we projected the template 3D volumes around the best viewing angles.

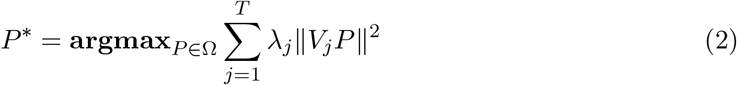

### 2.2 Linear Combinations of Template Conformations

In general, the success of matching experimental 2D-images to template 2D images depends on the accuracy of the chosen distance function: i.e. the two images should be highly similar when the pairwise distance between them is small. The distance is typically defined as the Euclidean distance of two images when they are optimally aligned. However, the alignment step is computationally expensive, this is because one image often needed to be rotated many times (*R* times) in order to obtain the best alignment with the other image. The cost was contributed mainly by the rotation operations and the frequent calculation of the Euclidean distances in the *N*^2^ × *N*^2^ dimensional space, resulting in a calculation complexity of *O*(*RM*^2^*N*^2^) with *M* being the number of images and *N* being the number of pixels. To reduce the computational cost, we developed an efficient two-stage method consisting of ACF and pixel-pixel matching, to match experimental 2D images with the template 2D images.

### 2.3 The first stage: matching based on ACF

The purpose of the first stage of matching is to efficiently remove the majority of the cryo-EM 2D images having viewing angles far away from the best viewing angles. Here, we adopted a new and efficient distance metric to characterize the difference between two images: the Euclidean distance defined by the low-dimensional ACF curves between two images. The ACF curve is a rotational and translational invariant representation of an image, which removes the need of alignment during distance calculation. The ACF curve was obtained as follows: Firstly, image binarization was performed using “random walk” to take image segmentation [17]. Then, the boundary points were extracted. Finally, we computed the autocorrelation (AC) of the elementary vectors, which connected the adjacent boundary points as shown in Figure 3(a). Each 2D image, either the template or cryo-EM 2D images, is then represented as an ACF curve, which is the plot of AC as a function of interval *n* (ACF curve in Figure 3(a). The AC at interval *n* is calculated by following equation (3):

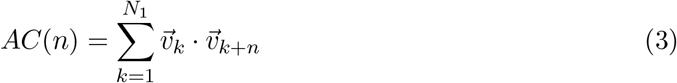

where 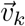 is the elementary vector connecting the *k_th_* boundary point and (*k* + 1)*_th_* boundary point and *N*_1_ is the number of boundary points.

**Figure 3:**
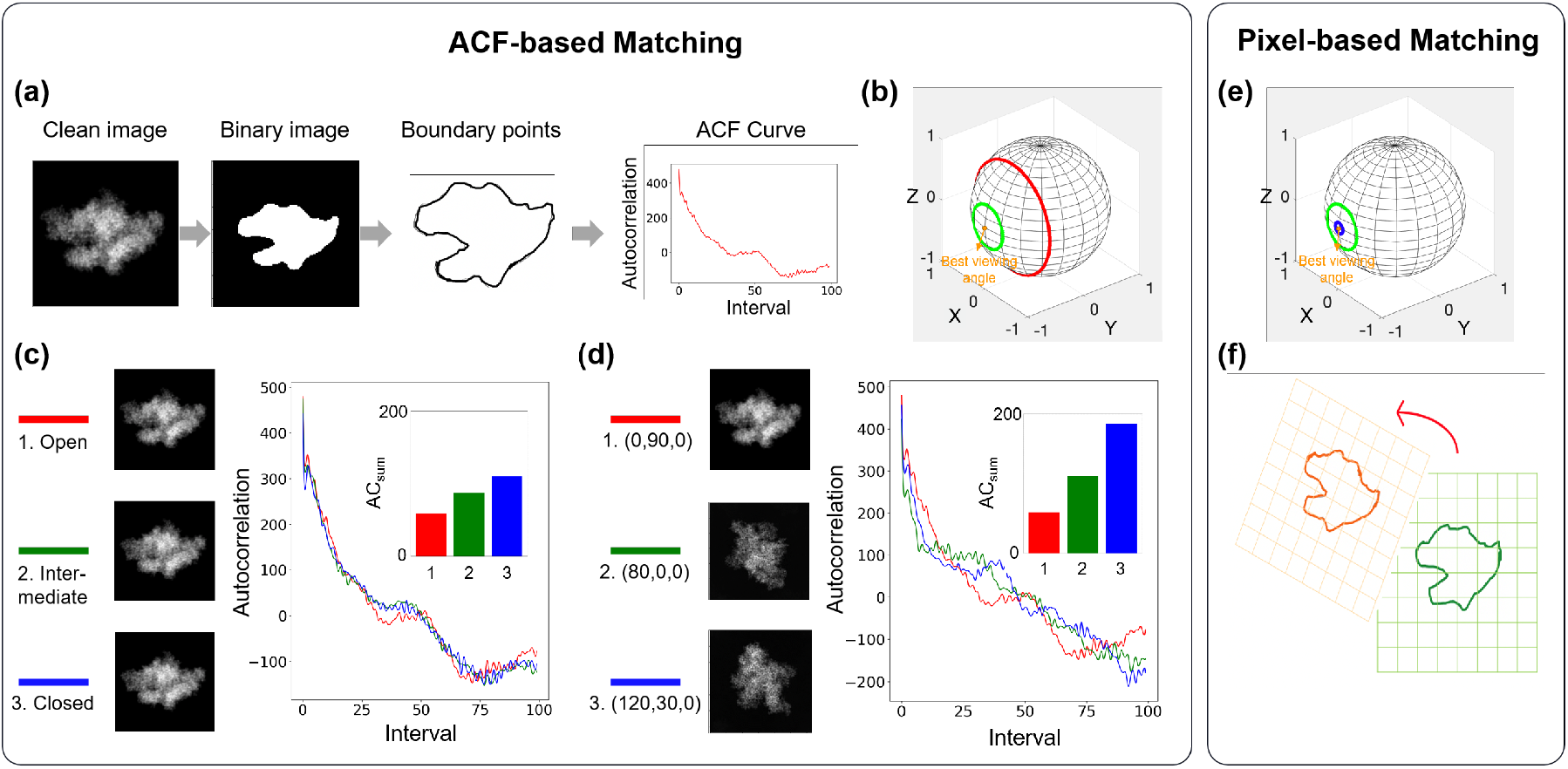
The two-stage matching of LCTC method. **ACF-based Matching** (a) In the first stage, the image was first binarized and the contour was extracted to generate the ACF curve. (b) The two ranges of viewing angles that were used to filter the cryo-EM 2D images projected far away from the best viewing angle. The middle and large range of viewing angles are shown as green and red circles. The best viewing angle is shown as orange dot. (c) The ACF curves are plotted for 2D images of 3D structures with different conformation projected on the same viewing angle. The inset bar-graph is the *AC_sum_* values. Both ACF curves and *AC_sum_* values of the 2D images of these 3D structures are similar. (d) The ACF curves and *AC_sum_* values of three 2D images of a 3D conformations projected on different viewing angles, which are very different from each other. **Pixel-based Matching**. The second stage is based on pixel-by-pixel matching. (e) The two ranges of viewing angles used for filtering the cryo-EM 2D images. The small and middle ranges are shown as blue and green circles. The best viewing angle is shown as orange dot. (f) Pixel-based matching depends on the calculation of distance based on all the pixel intensities of two images.

The distance calculation based on ACF is less computationally demanding as it depends on the number of boundary points, *N*_1_, which is fewer than the total number of pixels, *N*. It also avoids the alignment process. Thus, the calculation complexity is 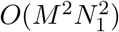, which is much less than the pixel-by-pixel distance calculation *O*(*RM*^2^*N*^2^). Although the calculation of ACF is computationally inexpensive, it can only distinguish the 2D images with different viewing angles and not the subtle conformational changes of the biomolecules. To demonstrate these points, we plot the ACF curves of the 2D images of a 3D conformation projected onto different viewing angles (Figure 3(d)) and calculate the summation of all AC values 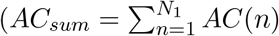, see the inset bar graph in Figure 3(d)). As shown by Figure 3(d), both ACF curves and *AC_sum_* are clearly different for these 2D images projected onto different viewing angles. On the other hand, as shown in Figure 3(c), the 2D images of three different 3D conformations (i.e. open, intermediate, and closed) projected at the same viewing angle produce similar ACF curve shape and *AC_sum_* values when projected onto the same viewing angle.

Therefore, we only use the ACF-based distance to filter out the cryo-EM 2D images having viewing angles far from the best viewing angle. To achieve that, we first defined two ranges of viewing angles around the best viewing angle, middle range (green circle) and large range (red circle excluding green circle) as shown in Figure 3(b). The template 3D conformations were projected onto the viewing angles in the middle and large range. Then, we calculated the pairwise Euclidean distance based on the ACF curve between the template images and all the cryoEM 2D images. The cryo-EM 2D images having distance closer to the middle range than the large range are kept, while the rest are filtered out. The ACF calculation is suitable for efficient comparison, yet it is less accurate since the ACF value only consider the boundary of the molecule neglecting the pixel intensity in the 2D images. Therefore, in the next stage, we used pixel-by-pixel euclidean distance calculation for accurate assignment of the cryo-EM 2D images to the template images.

### 2.4 The second stage: matching based on pixel-pixel

In the second stage, the purpose is to accurately assign the remaining cryo-EM 2D images from the first stage to the template images and estimate population of the template images. To ensure accurate assignment, we used distance metric based on the pixel-pixel intensities, which considers all the pixel intensities between two images after alignment. The alignment process requires frequent rotation and calculation of distance to find the optimum alignment between two images, which is computationally expensive. However, the comparison is done between the template images and a subset of the cryo-EM 2D images (instead of all of them), which alleviate the computational burden.

Furthermore, accuracy is also achieved by performing assignment only using the cryo-EM 2D images with viewing angles close to the best viewing angle, which can best distinguish the conformational changes of the biomolecule. Therefore to select the cryo-EM 2D-images projected near the best viewing angle, we defined two ranges of viewing angles, small and middle range (shown as blue and green circles, respectively in Figure 3(e)). Similar to the first stage, we projected the template 3D structures using the viewing angles in these two ranges. Then, we compare these template 2D images with the remaining cryo-EM 2D images from the first stage using the pairwise-distance based on pixel-pixel intensities. The cryo-EM 2D images having distance closer to the small range compared to the middle range are kept for assignment to template 2D images (shown in Figure 3(e)), while the rest are removed. Lastly, we counted the number of cryo-EM 2D images assigned to each template image within the small range, which reflects the population of the template images.

### 2.5 Comparison of LCTC with Maximum Likelihood-based and Clustering-based methods

We evaluated our LCTC method by comparison with the other two approaches: Maximum Likelihood-based method (add citation) and Clustering-based method.

To perform the Clustering-based method, we used manifold learning method to cluster the 2D images directly inspired by [14]. We first chose the best viewing angle, which can separate the three conformations in the *Taq* RNAP and *Eco* RNAP datasets separately. Then, the three template 3D volumes were projected onto a number of 2D images around the best viewing angle. Next, manifold learning was performed to extract the low-dimension feature, followed by K-means clustering to categorize the 30000 randomly chosen 2D images into three template conformations to determine their populations for both *Taq* RNAP and *Eco* RNAP datasets. For the Maximum Likelihood-based method, we performed Multi-body Refinement and then K-means clustering of their PCA eigenvectors to assign the cryo-EM 2D images into three groups.

In summary, we first validated the LCTC method and evaluated it by using a simulated dataset of RNAP clamp motion, where the ground truth is known. It is also demonstrated that the LCTC method outperforms the other two methods in terms of accuracy, as it can reproduce the populations of three *Taq* RNAP conformations. Furthermore, we successfully applied the LCTC approach in an experimental *Eco* RNAP cryo-EM dataset by revealing the populations of various clamp-opening conformations.

### 2.6 Simulated cryo-EM Datasets of *Taq* RNAP

Our simulated dataset was *Taq* RNAP system in complex with *σ_A_* factor, which is also called holoenzyme. To obtain the simulated dataset, we started with two crystal structures of *Taq* RNAP, which vary in their clamp conformation, open (PDB ID: 5TJG [18]) and closed (PDB ID: 4XLN [19]) clamp. For the closed clamp structure, the DNA and RNA were removed from the crystal structure, leaving only the RNAP and *σ^A^* for our dataset. We then generated the clamp intermediate structures between the open and closed clamps using multiple-basin coarse-grained (CG) molecular dynamic (MD) simulations ([20, 21]). From the initial CG-MD simulations, we chose three representative structures based on the clamp opening angle: open (12.3°), intermediate (8.4°), and closed (0°). These three structures (open, intermediate and closed) are back-mapped from coarse-grained to all-atom structures ([22]). We projected these three initial structures in a fixed proportion (gold standard) as our input 2D images and performed Multi-body Refinement analysis to reconstruct the three structures as templates. We chose clamp as the flexible part and the remainder of RNAP as the rigid part as the mask for Multi-body Refinement. We also randomly chose 30000 input 2D images, which would be assigned back to the three structures to determine their populations.

### 2.7 Experimental cryo-EM datasets of *Eco* RNAP

In *E. coli*, the essential primary *σ* factor, *σ*^70^, binds to RNA polymerase (RNAP) to form the *σ*^70^-holoenzyme that is capable of recognizing promoters and initiating the transcription of most genes in *E. coli*. We then applied LCTC to the published cryo-EM dataset of this complex. The 2D particle image is obtained from the EM databank (EMDataResource: EMD-20230). The pre-processing steps followed the original publication [23]. In the original publication, Multi-body Refinement, as implemented in RELION ([11]), was performed on the 370965 2D particle images to identify various clamp domain motions and the first three principal components (PCs) were reported. Moreover, we use RELION to reconstruct the 3D volume, then regard the projections as our denoised images. In this study, we focused on the second PC (PC2) corresponding to the Clamp’s twisting motion, as it has the largest variance of the Clamp twisting-open angle (*≈* 4.5°) [23]. The 2D images were classified into three classes based on PC2 and the 3D conformation of each class was reconstructed. We used these three reconstructed 3D volumes as our template dataset. Then, we randomly projected the three 3D volumes to 30000 2D images as our dataset.

## 3 Results and Discussions

### 3.1 Application of LCTC to the simulated *Taq* RNAP dataset

To validate our method, we created a simulated dataset of *Taq* RNAP in complex with *σ* factor with different degrees of the clamp domain opening (yellow domain in Figure 2(b),(c),(d)). This complex binds to the promoter DNA during the initiation of transcription and subsequently loads the DNA into the RNAP active site. The clamp domain of RNAP acts as a gate to allow the DNA to pass through, thus the opening and closing of the clamp domain are critical during DNA loading.

To search for the best viewing angle with the largest variance that can distinguish the clamp opening, we randomly chose 2562 viewing angles in the sphere. To quickly calculate the variance of different conformations in each viewing angle, we first performed PCA of the whole three 3D volumes of template structures reconstructed by Multi-body Refinement to reduce dimension. We chose the first eigenvalue according to Figure 4(a). Then, the best viewing angle was identified as (*φ, θ, ψ* = 0, 90, 0). The variances of the three conformations in different viewing angles are plotted as – log(Var) (Figure 4(b)), which shows that the (0,90,0) has the largest variance.

**Figure 4:**
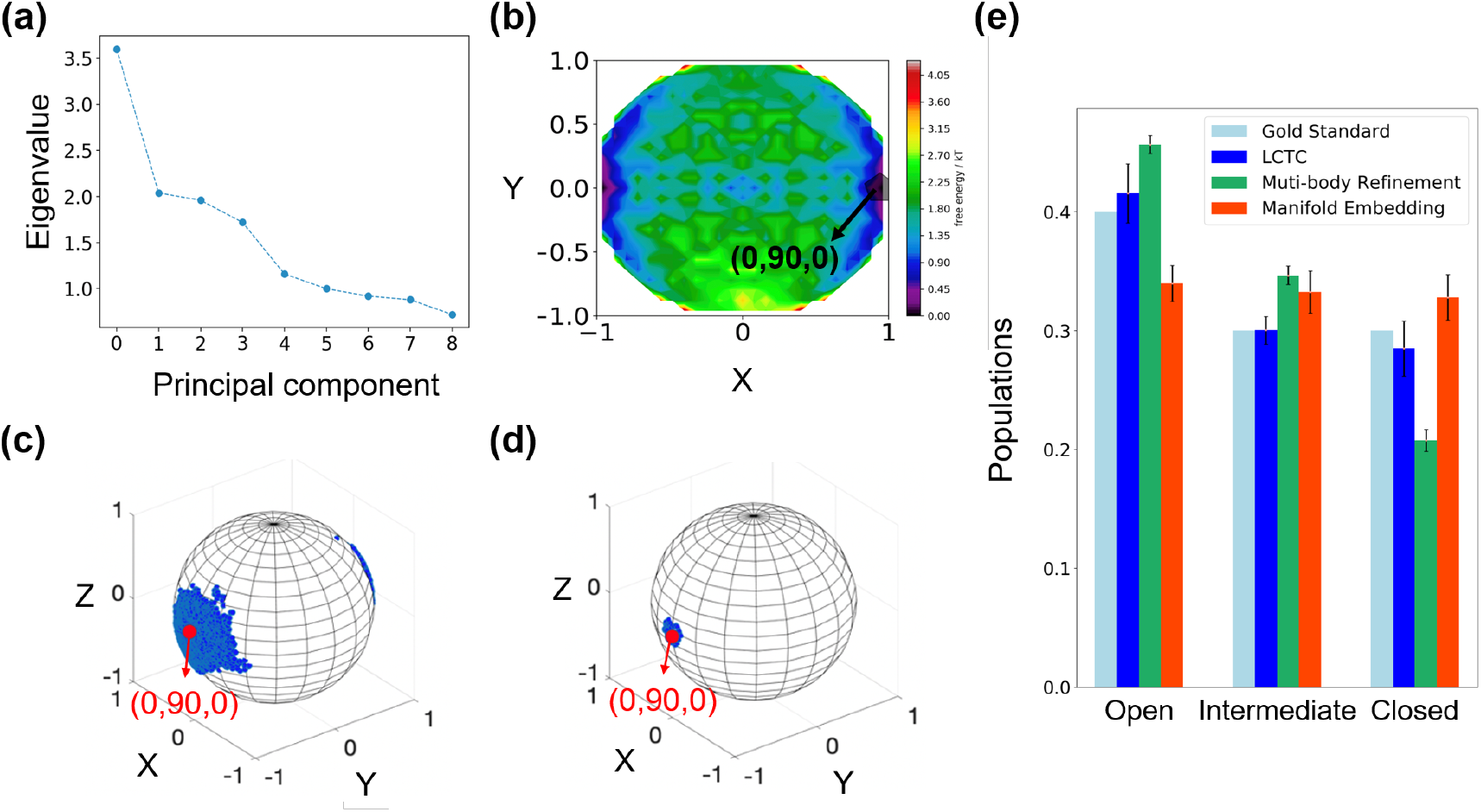
Application of LCTC on the simulated dataset of Taq RNAP. (a) Eigenvalues obtained from PCA of the template 3D volumes were plotted. (b) The variance of three conformation distribution in the sphere. Projecting from the north pole, the darker color represents higher variance. (c) The viewing angles of the remaining cryo-EM 2D images after ACF matching were shown as blue points. (d) The viewing angles of the cryo-EM 2D images used for assignment after pixel-pixel matching were shown as blue points. The best viewing angle is shown as red dot in Figure (c) and (d). (e) The populations of the three template conformations calculated by different methods: LCTC method, Clustering-based method, Maximum Likelihood-based method, and the gold standard.

Next, we randomly selected 30000 cryo-EM 2D images from the cryo-EM dataset to assign to the three template structures. Furthermore, we projected the template structures around the best viewing angle. To perform the two-stage matching, we defined three ranges of viewing angles around the best viewing angle. The small, middle, and large ranges cover the spherical cap of 5, 20, 60 degrees cone angle around the best viewing angle, respectively. The large range includes the Euler angles (*φ, θ, ψ*) (*φ* = 0, 30, 60, …, 330; *θ* = 30, 40.., 90; *ψ* = 0), with a total of 84 viewing angles. The middle range includes (*φ, θ, ψ*) (*φ* = 0, 30, 60, …, 330; *θ* = 10, 15, 20; *ψ* = 0), with a total of 36 viewing angles. The small range includes (*φ, θ, ψ*) (*φ* = 0, 30, 60, …, 330; *θ* = 0, 2.5, 5; *ψ* = 0), with a total of 25 viewing angles. In the first stage, we collected the 2D images having viewing angles within the middle range (Figure 3 (b)), regardless of the clamp conformations of RNAP. We extracted 100 boundary points for the calculation of ACF curves. As the simulated dataset contained clean projections, we binarized the images by setting the signals on the image as 1 and the backgrounds as 0. As shown in Figure 4(c), the blue points represent the viewing angles of the remaining 2D images after the first-stage of matching. This result clearly showed that the ACF-based matching can separate images from different viewing angles. The ACF stage matching has successfully removed about 90% of the dataset, hence only about 3000 images were used for the second stage of matching with higher accuracy.

In the second stage, we assigned the 2D images that are projected within the small range to the template conformations (Figure 3(e)). Then, we counted the 2D images assigned to each template image and obtained the populations by distributing these images to three template conformations. There are around 300 images remaining having viewing angle within the small range for the assignment and their viewing angles are shown in Figure 4(d).

The population distribution is shown in Figure 4(e). For the test data, we set the initial proportions as 0.4, 0.3, 0.3 to be the gold standard for open, intermediate and close conformation and performed three trials of the experiment. Importantly, our calculated results from LCTC were consistent with the gold standard. In contrast, Manifold Embedding and Multi-body Refinement method generated the gold standard that lay outside the populations. We speculated that the Manifold Embedding method obscured the minor difference of multiple conformations because some information may be lost due to the process of reducing dimension. Also, the populations obtained directly from 3D volumes that was recovered from Multi-body Refinement cannot reproduce the gold standard. Therefore, the LCTC method is the best method to distinguish the conformational change in simulated *Taq* RNAP dataset.

The Taq RNAP Holoenzyme dataset studies in the paper was a simulated dataset in the absence of the noise. In practice, the difference between various conformations in a structural ensemble may be obscured by the noise. Therefore, we have performed an experiment using noisy cryo-EM 2D images of the simulated *Taq* RNAP dataset. We showed that even with signal to noise ratio (SNR)=0.1 of the cryo-EM 2D images, we can reproduce the population of the template 3D conformations accurately (Appendix 6.2).

### 3.2 Application of LCTC to the *Eco* RNAP

Similar to *Taq* RNAP, the complex of *Eco* RNAP and *σ*^70^ factor binds to the promoter DNA and opening of the clamp domain facilitates the loading of the DNA into RNAP. The degree of the clamp opening can significantly affect the DNA loading process. Specifically, the binding of a transcription factor TraR was shown to limit the movement of the clamp domain, which in turn either hinder or activate DNA loading [23]. Thus, accurate prediction of the populations of Clamp domain conformations is important to determine the degree of Clamp opening.

After confirming that our algorithm performed well in the simulated *Taq* RNAP dataset, we applied our algorithm in the experimental *Eco* RNAP dataset. The procedures used for the experimental data were mostly the same as simulated data. The experimental dataset consists of *Eco σ*^70^-holoenzyme, which is a complex of *Eco* RNAP and *σ*^70^. We firstly pre-processed the cryo-EM data by RELION: we used RELION to denoise the noisy data, which consists a total of 370965 2D images as shown in Figure 5(a). The reconstructed 3D volumes from Multi-body Refinement performed on the clamp domain were used as template structures [23] (shown in Figure 5(c)). Next, we selected the best viewing angles (0, 0, 0) (shown in Figure 5(b)) based on the three template structures’ eigenvectors. The three ranges of the viewing angles were the same as the simulated dataset. Finally, the results of the population are shown in Figure 5(d). As shown before, our algorithm matched the best to the gold standard in comparison with other methods. Thus, it is implied that the populations calculated by LCTC may reflect the native equilibrium population of the *Eco* RNAP clamp domain. Importantly, we have demonstrated that the LCTC method can elucidate the dynamic ensemble of protein conformations from experimental cryo-EM datasets.

**Figure 5:**
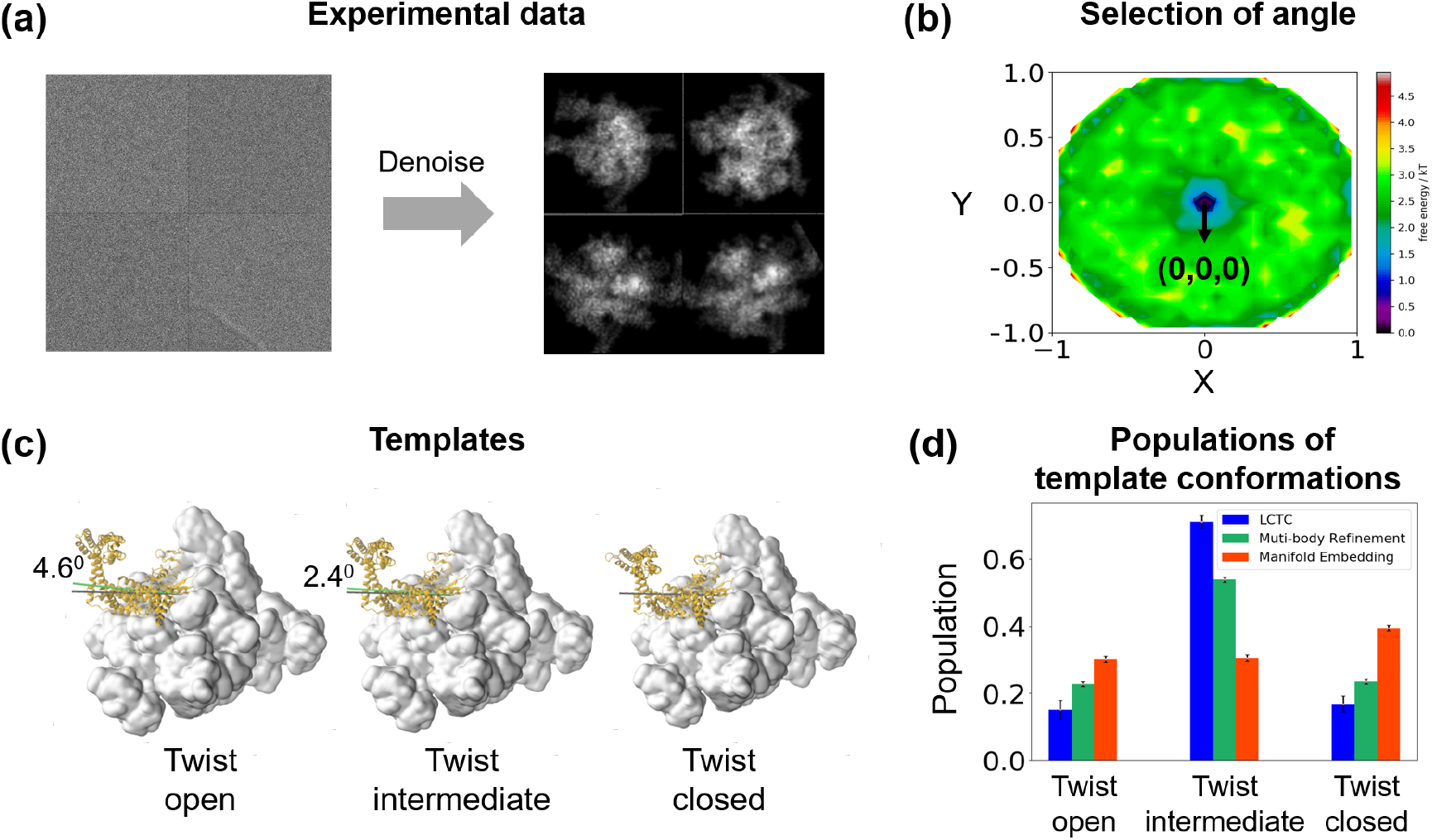
Application of LCTC on the experimental dataset of Eco RNAP: (a) The generation of the denoised 2D images as input data. (b) The selection of the best viewing angles, (0,0,0). (c) The three template conformations obtained by Multi-body Refinement (d) The populations of three template conformations obtained by different methods: LCTC, Multibody Refinement, and Manifold Embedding.

With the current advancement of cryo-EM technology, multiple classes of biomolecule conformation can be identified from a single biological sample. These classes of conformations that are present in the solution may be functionally relevant to the biological pathway of the biomolecule. An algorithm to reconstruct the free energy landscape of a biomolecule has recently been developed, which is based on manifold embedding of the cryo-EM 2D images [24], and obtain the functional conformational changes of the molecule. Interestingly, a diffusion map-based algorithm has been developed to elucidate the biological pathway based on populations of intermediate conformations, with the augment of molecular dynamics simulations [25]. In combination with these two methods, we anticipate that our LCTC method holds the promise to be applied to elucidate mechanisms of functional conformational changes in the future. In addition to the Multi-body Refinement on experimental dataset, Markov State Model analysis of molecular dynamics simulations provides an alternative approach to generate metastable intermediate conformations as template 3D structures in our LCTC method[26, 27, 28].

## 4 Conclusion

In this work, we have presented an LCTC algorithm to identify the population distribution of the multiple conformations existing in the heterogeneous cryo-EM dataset. The idea is to use template 3D conformations, obtained by Multibody refinement, and assign the cryo-EM 2D images to these template structures. We first identify the best viewing angle that can best distinguish the conformational changes. Then, the cryo-EM 2D images are assigned to the template images projected at the best viewing angles. To assign the cryo-EM 2D images efficiently, we use the new distance metric based on ACF curve to remove the majority of irrelevant 2D images. Then, the accurate assignment is done using the pixel-based matching between the remaining 2D images with the template images projected at the best viewing angle. Furthermore, the successful application to the RNAP dataset demonstrates the potential utility of our method in identifying the population distribution of conformations in the experimental cryo-EM images, and thus may help elucidate the underlying free energy landscape of macromolecules.

## 5 Acknowledgement

We thank helpful discussions from Dr. Michael Levitt. We also thank Dr. Seth Darst for providing the Experimental cryo-EM datasets of *Eco* RNAP. X.H. acknowledges support from the Padma Harilela endowment fund. The research of Y.Y. was supported in part by Hong Kong Research Grant Council (HKRGC) grant 16303817, ITF UIM/390, as well as awards from Tencent AI Lab, Si Family Foundation, and Microsoft Research-Asia.

## 6 Appendix

## 6.1 The explanation of using PCA to calculate variances in multiple conformations

In the Section 2.1, we took PCA of *K* representative conformations firstly to simplify the computing and calculate the variance of *K* conformations based on principal component values. The following Equations (4) and (5) prove the calculation of variances through top *T* eigenvectors approximates the estimate of variances directly, where *P* is projection operator represented as a *N*^3^ × *N*^3^ matrix related to the viewing angle, λ_1_, λ_2_… are eigenvalues in descending order, *A* is a large 3D volume matrix consisting of *K* conformations *A_i_* (*i* = 1, 2, …*K*), Σ is the covariance matrix, and *V_j_* is the *j*-th row of *V*.

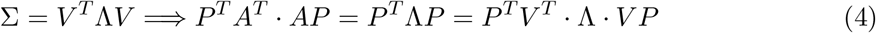

which implies that

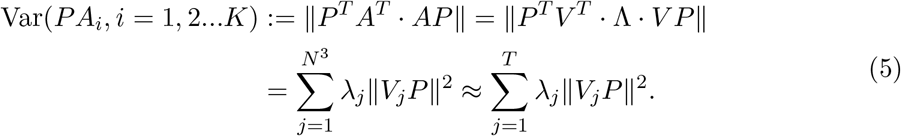

## 6.2 Robustness for simulated noise of LCTC

In order to test the robustness for noise of our method, we generate the simulated dataset with Gaussian noise. Specifically, we generate some random projections for three conformations of simulated cryo-EM datasets of Taq RNAP. We contaminate those clean images with additive Gaussian noise at different SNR (0.1 and 0.05) where the SNR is defined by “SNR =Var(Signal)/Var(Noise)” in the real space. For simplicity, we did not apply the contrast transfer function (CTF) to the datasets, and all the images are centered.

We borrowed from [29] as our denoising method, which is inspired by the *β*-GAN [30]. Then we set the initial proportion of open, intermediate and closed conformation as 0.4, 0.3, 0.3. After denoising, the population results obtained by our LCTC method are shown in Figure. It demonstrates that LCTC method performs well in accurately reproducing the populations of three RNAP structures with different degrees of the clamp opening even the data is contaminated with the simulated noise.

**Figure 6:**
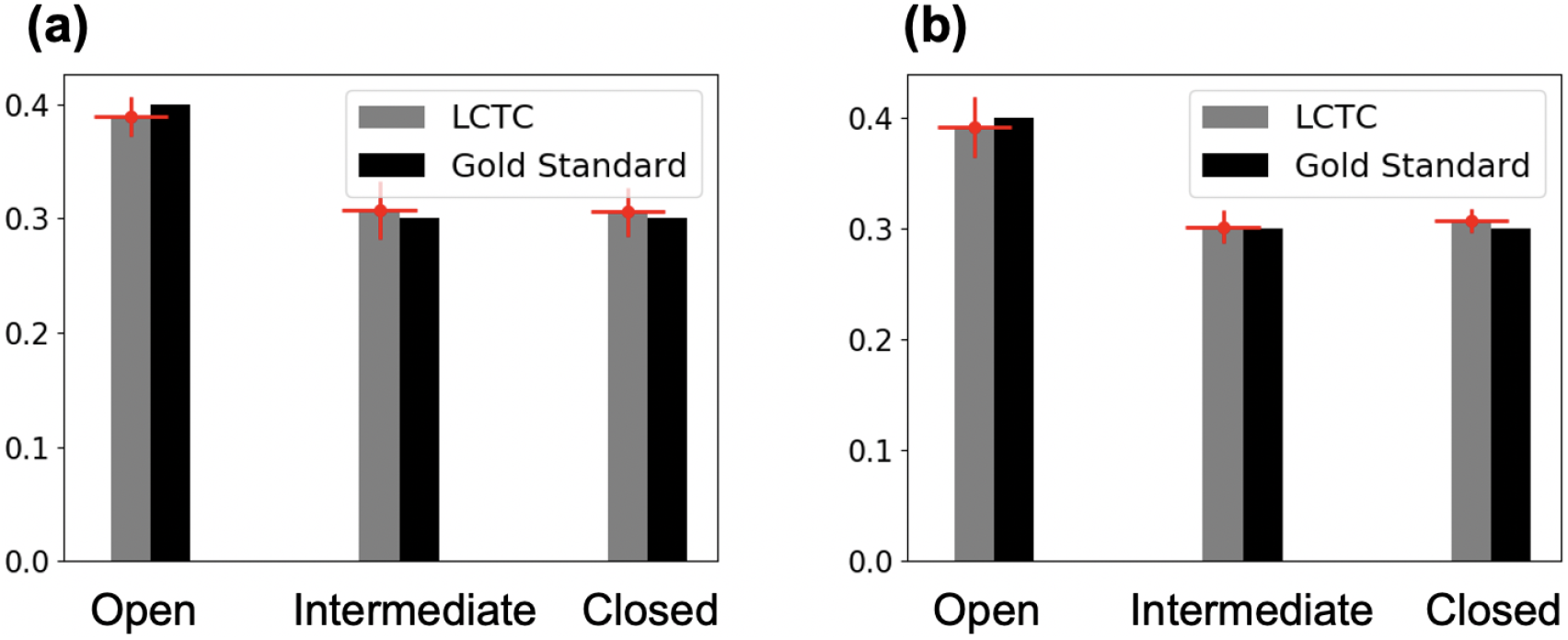
The population results in Taq RNAP data with SNR 0.1 and 0.05. (a) Population result in SNR 0.1. (b) Population result in SNR 0.05.

